# CobVar — a comprehensive resource of Vitamin B_12_-associated genomic variants

**DOI:** 10.1101/2025.02.05.636608

**Authors:** Neha HS, Chaitra A Kilpady, Akshata A Kamath, Ashikh Ahmed, Tinku Thomas, Anura V Kurpad, Ambily Sivadas

## Abstract

The importance of vitamin B_12_ (cobalamin) in numerous biological processes, including DNA synthesis and cellular energy production, underscores the need for therapeutic and public health strategies to address B_12_ insufficiency/deficiency in the population. Genetic variations in pathways influencing cobalamin absorption, transport, and metabolism can affect various direct and indirect measures of vitamin B_12_ status. To facilitate a structured approach to studying these genetic factors, we aimed to systematically curate and create a user-friendly web database that offers comprehensive data on genetic variants influencing B_12_ biomarkers. A Pubmed search was performed for 5 B_12_ traits (total serum/plasma B_12_, holotranscobalamin (active B_12_), total transcobalamin, holo haptocorrin, and methylmalonic acid) resulting in 493 research publications, of which 47 relevant publications were reviewed further. The database backend was built using MongoDB and the web interface was coded in PHP, JavaScript, HTML and CSS on an Apache HTTP server. We have manually curated and compiled the Cobalamin Associated Genetic Variant (CobVar) database, comprising a total of 324 genetic variant associations for 5 different vitamin B_12_ traits involving 222 unique genetic variants and 84 genes identified across several genome-wide association studies (GWAS) and candidate gene studies. About one-third of the total genetic variant associations have been reported in >1 independent studies and 15 variants in >1 ethnic group. FUT2 gene showed the maximum number of associations for total serum/plasma B_12_ (N=39), followed by MTHFR (N=24) and TCN2 (N=23). The database is accessible online at https://datatools.sjri.res.in/VBG/. CobVar is a vital resource for researchers and nutritionists, offering quick access to the latest developments in B_12_-related genetic variant research and serves as a valuable tool for advancing personalized treatment.

**Database URL:** https://datatools.sjri.res.in/VBG/.

## Introduction

Cobalamin, commonly known as vitamin B_12_, is an indispensable water-soluble micronutrient, primarily found in animal food sources. It is essential for various physiological processes, including normal functioning of the nervous system and the development and maturation of red blood cells. It serves as a cofactor for two critical enzymes: cytosolic methionine synthase, necessary for nucleic acid synthesis through the regeneration of tetrahydrofolate, and mitochondrial methylmalonyl-CoA mutase, which supports the citric acid cycle and heme synthesis(1,2). It also plays key roles in DNA synthesis, the methylation cycle, and energy production, and is crucial for maintaining cell membranes and the myelin sheath, which protects nerves and ensures proper transmission. Vitamin B_12_ deficiency can manifest in both clinical and sub-clinical forms, with significant health implications. Clinical B_12_ deficiency (usually defined as a total serum B_12_ of <150 pmol/L) may manifest as neuropathy, cognitive impairment, and anemia(3), while sub-clinical deficiency (usually defined as a total serum B_12_ of <200 pmol/L), though asymptomatic, is equally important to detect as it can still cause metabolic disruptions over time(4).

Assessing vitamin B_12_ status involves several biomarkers that provide insight into both circulating levels and the functional impact of deficiency. Total circulating B_12_ is a commonly used measure that reflects the overall concentration of vitamin B_12_ in the blood, though it may not always accurately reflect tissue availability or early deficiency(5). Holotranscobalamin (holoTC), the biologically active form of B_12_ bound to transcobalamin, offers a more precise indicator of B_12_ available for cellular uptake, making it a sensitive marker for detecting early deficiency. Holohaptocorrin, another form of B_12_ bound to haptocorrin, represents a storage form of B_12_ that is not readily available to cells, and its clinical utility is less direct compared to holoTC. Functional biomarkers like methylmalonic acid (MMA) and total homocysteine (tHcy) accumulate when B_12_ levels are insufficient. Elevated MMA is a specific marker for B_12_ deficiency, as B_12_ is required for its metabolism. Total homocysteine (tHcy), while indicative of B_12_ deficiency, can also be elevated due to low levels of folate, riboflavin, or vitamin B6. These functional biomarkers are especially useful for detecting subclinical B_12_ deficiency and early changes in B_12_ status. Together, these biomarkers provide a comprehensive evaluation of an individual’s vitamin B_12_ status.

Vitamin B_12_ biomarkers exhibit significant inter-individual variability, influenced by several factors, including age, diet, gastrointestinal health, pregnancy, other comorbidities and medications, and genetic factors(6). Studies show that 59% of the variability in plasma vitamin B_12_ levels can be attributable to genetic factors(7). Over the years, numerous candidate gene studies and genome-wide association studies (GWAS) have identified multiple genetic markers linked to vitamin B_12_ status. Candidate gene studies, which focus on specific genes involved in B_12_ pathways, have uncovered polymorphisms in genes such as FUT2, TCN2, and MTHFR, all of which play crucial roles in B_12_ absorption, distribution, metabolism/utilization, and excretion [ADME](8–10). More recently, large-scale GWAS have expanded this understanding by scanning the entire genome, leading to the identification of several novel genetic loci associated with various B_12_ biomarkers(11).

Despite the progress made through these diverse and isolated studies, there remains a significant gap in consolidating this information into a unified resource. Currently, there is no comprehensive, curated database that compiles all known genetic variants associated with vitamin B_12_ status and biomarkers, along with evidence of replication across multiple studies and ethnic groups. This lack of integration poses a challenge for efficiently accessing and interpreting genetic data, limiting its potential application in precision nutrition and clinical care. To address this, we aimed to systematically curate all B_12_-associated genetic variants reported in the literature and develop an online centralized database, making this valuable information easily accessible for researchers, nutritionists, and clinicians alike.

## Materials and methods

### Data curation from published peer reviewed literature

We searched in PubMed (http://www.ncbi.nlm.nih.gov/pubmed) and identified 493 original research publications (as of 25^th^ August 2024) for curation of B_12_ associated genetic variants. The search string utilized a combination of MeSH Terms and plain text of relevant keywords - (humans[MeSH Terms]) AND ((“vitamin b 12/blood”[MeSH Terms] OR Vitamin B 12 / metabolism[MeSH Terms] OR Vitamin B 12 Deficiency / genetics[MeSH Terms] OR Vitamin B 12 Deficiency / blood[MeSH Terms]) AND ((association study, genome wide[MeSH Terms] OR Genetic Association Studies[MeSH Terms]) OR ((polymorphism, single nucleotide[MeSH Terms] OR Polymorphism, Genetic[MeSH Terms]) AND (genotype[MeSH Terms])))) OR ((“Vitamin B12”[Title] OR B12[Title] OR Cobalamin[Title] OR homocysteine[Title]) AND (vitamin B12 [Title/Abstract] OR cobalamin[Title/Abstract]) AND (Allele[Title/Abstract] OR Polymorphism[Title/Abstract] OR Polymorphisms[Title/Abstract])) NOT (review[Publication Type])

After manual inspection of the 493 publications, 47 relevant publications were systematically reviewed to collate the reported genetic variant associations with 5 B_12_-related traits (total serum/plasma B_12_, HoloTC, HoloHC, MMA and Total TC). Various parameters from the publications including the method of study, study design and geographical location, ethnicity, etc. were documented in a pre-formatted tabular template.

### Database and interface design

The backend architecture of the database (CobVar) is built using open source web technologies like MongoDB, PHP, and JavaScript and the server was hosted using Apache HTTP server, creating a robust system for managing and querying large datasets of genetic variants. MongoDB serves as the primary database, allowing for flexible data storage across various fields related to genetic associations, such as ‘dbSNP ID’, ‘Gene’, and ‘Phenotype/Trait’. PHP facilitates server-side processing, handling requests, executing queries, and dynamically generating responses based on user input. This integration enables efficient data retrieval and ensures that users can access relevant information swiftly.

On the client side, JavaScript enhances user interaction and experience by enabling dynamic features such as real-time updates and interactive charts. The use of AJAX further optimizes performance, providing an efficient way to load data asynchronously while maintaining a responsive interface.

## Results

### Data curation

The database catalogs a total of. The data was systematically curated from a total of 47 peer-reviewed publications including, 34 candidate gene and 13 genome-wide association studies (GWAS) (**Figure 1**).

**Figure 1.**
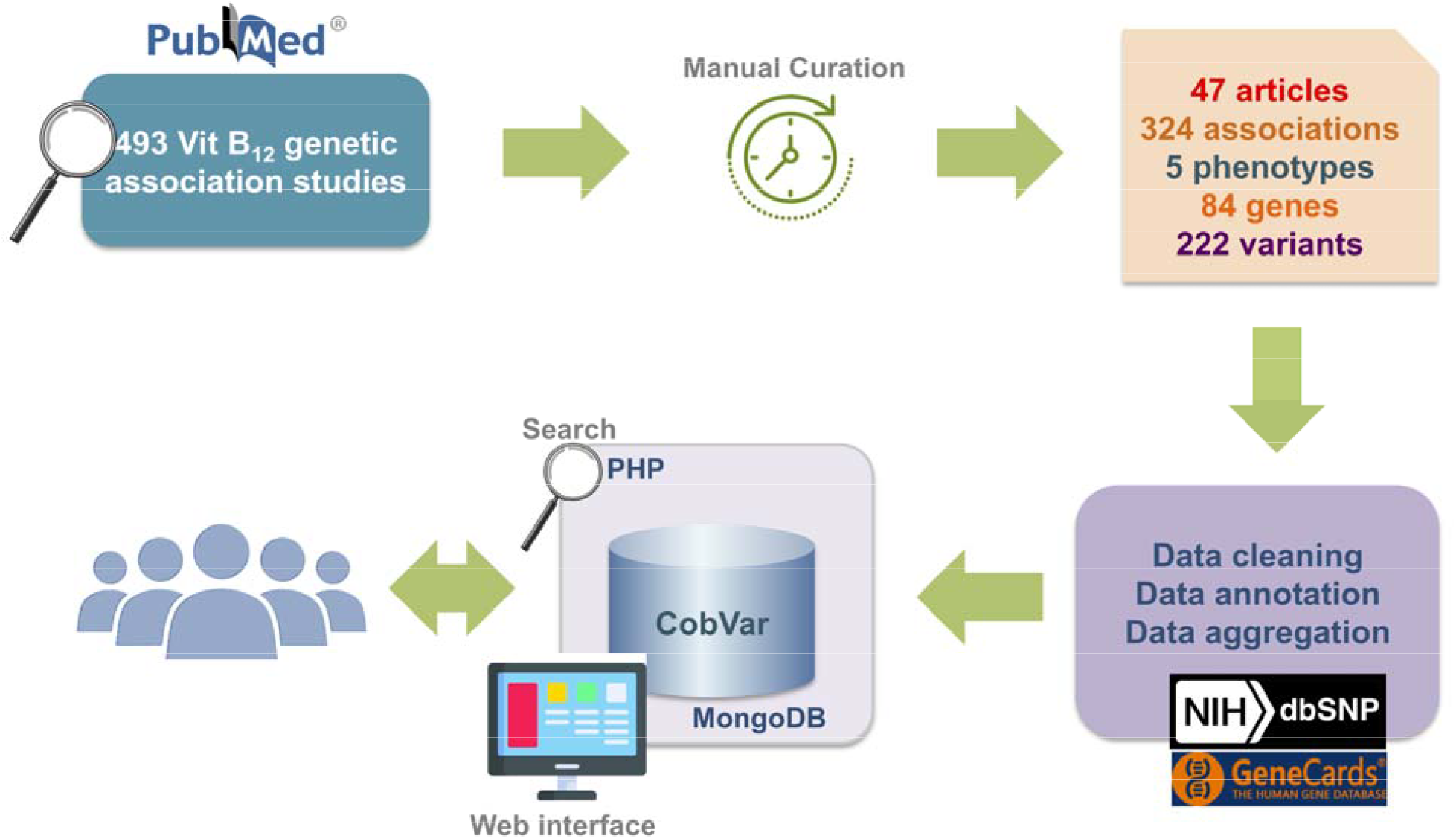
A schematic workflow of data collection and database integration for CobVar database

The studies were conducted primarily across 5 ethnic groups, with European (which included ethnicities like Irish, Icelandic, Danish, Greek, Norwegian and Canadian) comprising the largest proportion (43%) followed by Brazilian and South Asia (which included Indian and Pakistani ethnicities). In our study, around 33.6% of the total genetic variant associations have been reported in >1 independent study. This includes 2 total B_12_ associations replicated in >10 independent studies (rs1801133: MTHFR; rs1801198: TCN2), and an additional 7 total B_12_ associations replicated in ≥5 studies (rs526934: TCN1; rs602662: FUT2; rs601338: FUT2; rs1805087: MTR; rs1801131: MTHFR; rs492602: MTHFR; rs1141321: MUT). Two variants, rs1801133 (MTHFR) and rs1805087 (MTR), were replicated in >3 ethnic populations, 7 variants in >2 ethnicities and 15 variants in >1 ethnicity (**Supplementary Table 1**).

The largest number of variants curated was (N=155) for circulating levels of total serum/plasma B_12_ (48%, **Figure 2**). HoloTC showed 68 associations followed by holoHC with 67 associations. Maximum variant overlap was observed for total B_12_ and holoHC.

**Figure 2.**
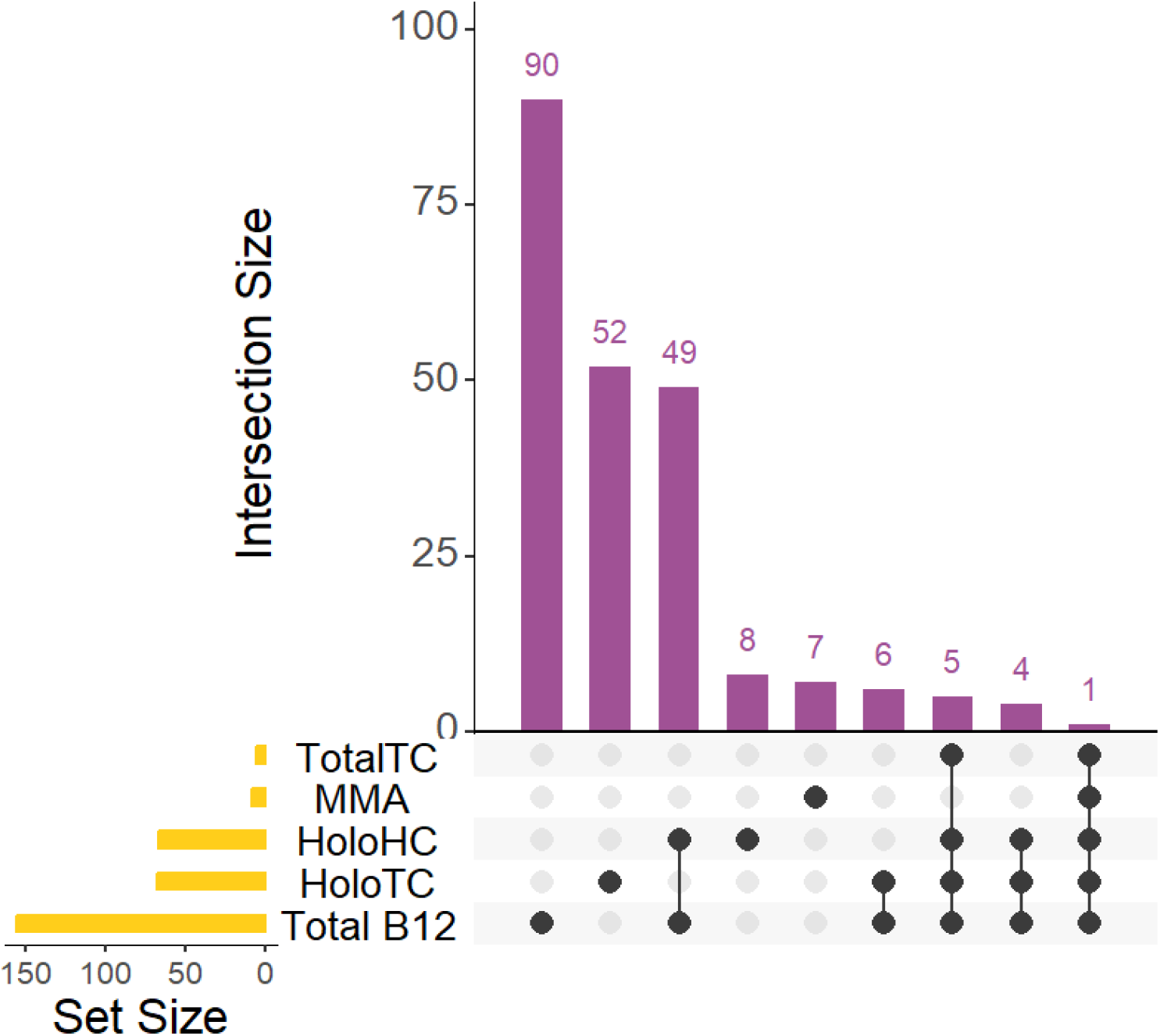
An overlap analysis of genetic variant associations across B_12_ traits.

Several genes involved in the absorption, distribution, metabolism/utilization and excretion pathways of vitamin B_12_ have been associated with different B_12_ traits. FUT2 gene showed the maximum number of associations for total serum/plasma B_12_, with 39 variants, followed by MTHFR (24 variants) and TCN2 (23 variants) (**Figure 3a**). Similarly, for holoTC, cubulin gene (CUBN) shows the maximum number of associations (14 variants) followed by TCN2 (12 variants) and CD320 (4 variants)(**Figure 3b)**. HoloHC showed maximum associations with FUT2 Gene (13 variants), followed by IZUMO1 (18 variants) and CUBN (5 variants)(**Figure 3c)**. MMA phenotype showed maximum associations with HIBCH (2 variants) while totalTC showed 1 variant association each in CUBN, FUT2, MMUT, MTR, TCN1 and TCN2 genes (**Figure 3d,3e)**.

**Figure 3.**
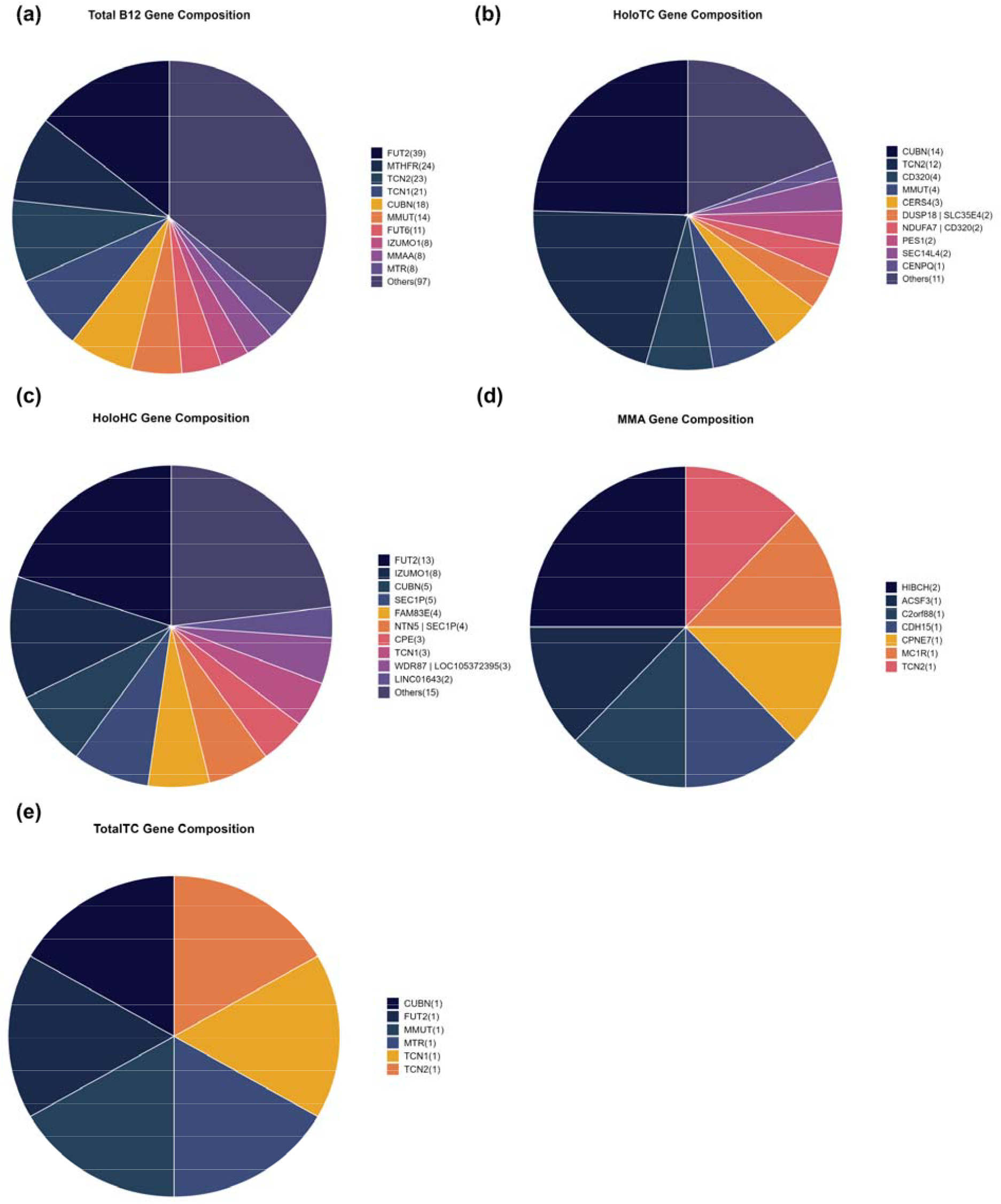
Gene distribution for different B_12_ trait associations. The number in parentheses next to each gene name represents the count of reported variants within that gene.

### Genetics of B_12_ homeostasis

The curation conducted in this study allows us to summarize the key genes involved in the complex process of vitamin B_12_ absorption, transport and metabolic utilization (**Figure 4**). TCN1, CUBN, and TCN2 are involved in the initial absorption and transport phases, ensuring that B_12_ reaches tissues where it’s needed. MMACHC, MTR, MTHFR, and MUT facilitate B_12_’s metabolic functions in DNA synthesis and mitochondrial energy production, while CD320 aids in cellular uptake. Methylation processes are also regulated by enzymes requiring B_12_ as a cofactor, impacting cellular functions. This complex network highlights the essential roles of these genes in maintaining B12 homeostasis and cellular function.

**Figure 4.**
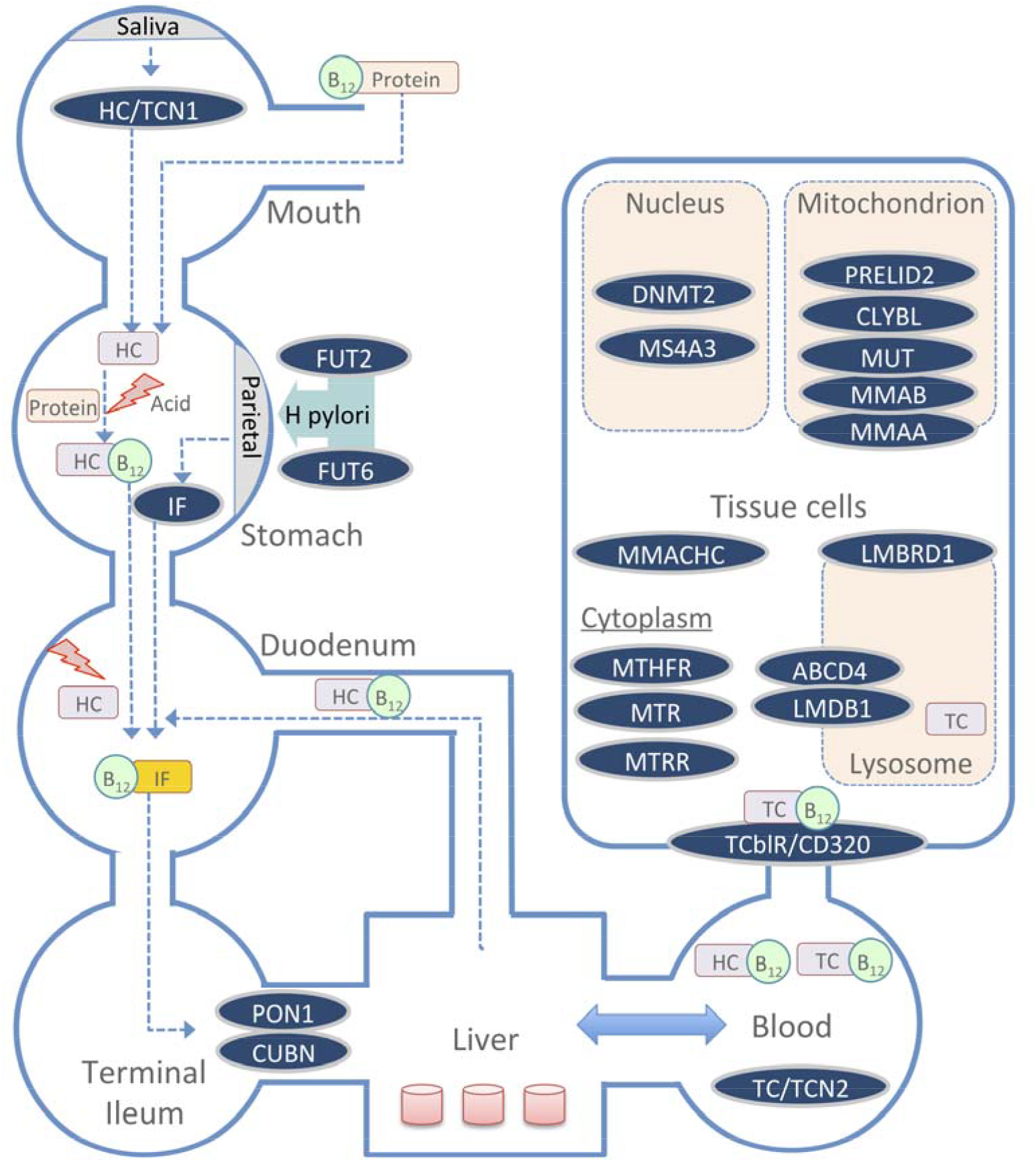
A summary schematic of the key genes involved in the absorption, transport and metabolic utilization of B_12_ across different body compartments.

### Database and web interface features

The database features a user-friendly interface that enables users to textually search based on various query terms, including dbSNP ID, chromsome-position, gene name, and specific B_12_ traits (**Figure 5**). The search results are displayed in a concise table of matching genetic variant associations, with complete information accessible by clicking on individual rows **(Figure 6a)**. The variant associations are organized into two main sections: the *Variant* Section **(Figure 6b)** and the *Association* Section **(Figure 6c)**. The *Variant* Section provides key annotations related to the genetic variant, with external links to databases such as dbSNP and GeneCards for further details. The *Association* Section presents a series of study results (referenced by PubMed IDs), displaying relevant association statistics and study details in a sequential format. The study details include ethnicity, geographical location, specific B_12_ trait and association statistics including effect size and p-values. The *Home* page includes an interactive pie chart summarizing the gene distribution across various trait associations, along with a brief overview of key data statistics. Additionally, a comprehensive user manual is available to guide users in navigating and effectively utilizing the database. The database can be freely accessed at https://datatools.sjri.res.in/VBG/

**Figure 5.**
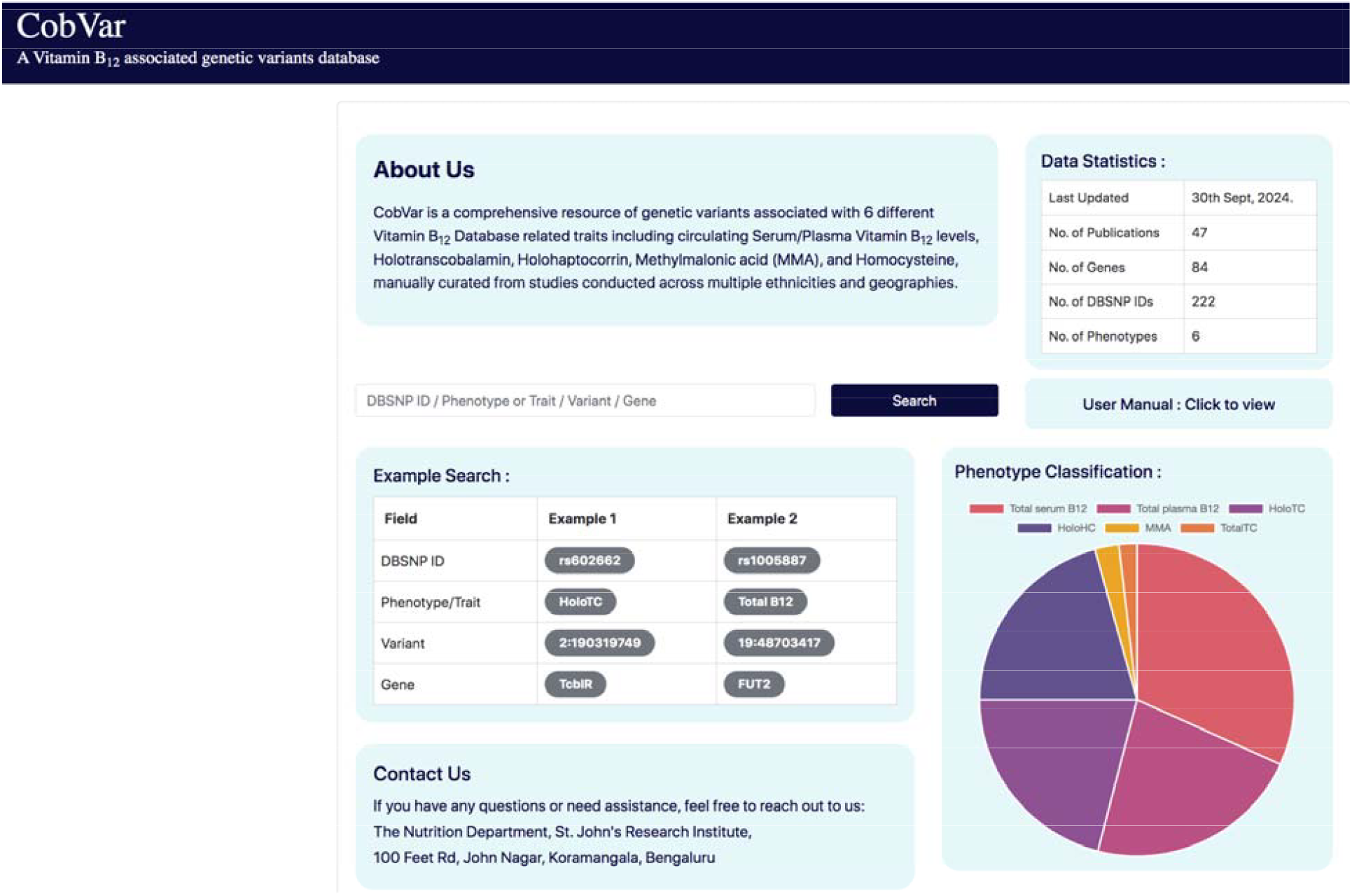
Screenshots of the CobVar database application featuring the Home page, and the different search query formats that can be used by the user.

**Figure 6.**
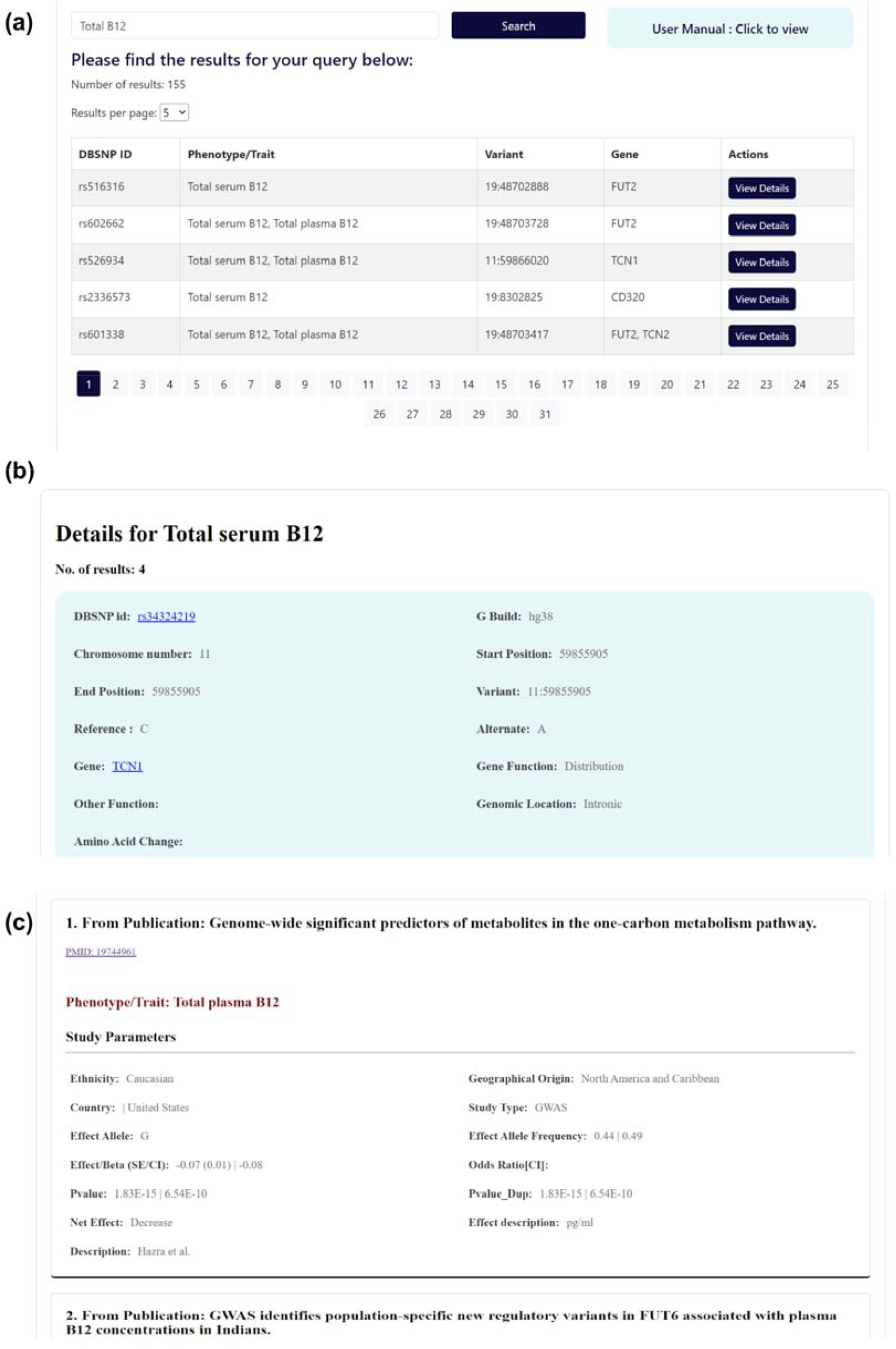
Screenshots of **(a)** the expanded database query results, **(b)** the detailed view of the *Variant* section and **(c)** the *Association* section.

## Discussion

The CobVar database offers a centralized platform that consolidates curated genetic data linked to B_12_ homeostasis, enabling a deeper understanding of the genetic factors influencing B_12_ absorption, transport, clearance and utilization. The database helps identify high-confidence variant associations that have been replicated across multiple studies and diverse populations, ensuring robust and reliable data for research and clinical use. The database can also be utilized to generate polygenic risk scores for populations by aggregating the impact of multiple genetic variants associated with vitamin B_12_ biomarkers, enabling more accurate assessments of genetic predispositions to B_12_ deficiencies/insufficiency and other related health outcomes, including anemia and neurological disorder.

The database will also enable nutritionists to develop personalized nutrition strategies by incorporating genetic data into dietary recommendations. As genetic testing becomes more accessible, understanding individual differences in B_12_ status can help tailor supplementation and dietary plans based on a person’s genetic profile. This approach can optimize B_12_ intake, ensuring that patients and clients receive the most effective nutritional advice, particularly those at risk for deficiency due to genetic factors. This database thus serves as a valuable tool for advancing both precision nutrition and genomic research.

## Conclusion

With emerging evidence on novel and population-specific genetic variants associated with various direct and indirect measurements of vitamin B_12_ status, there is increased scope for exploring the physiological and molecular underpinnings of the ADME pathways associated with vitamin B_12_ in humans. These findings carry important consequences for understanding the genetic contributors to B_12_ deficiency and tailoring treatment strategies. To keep up with the rapid advancements in this field, the CobVar database will be updated periodically. These updates will incorporate data from both peer-reviewed studies and preprints, ensuring comprehensive and up-to-date annotations. This resource is poised to become a vital hub for researchers and nutritionists, offering free and quick access to the latest developments in B_12_-related genetic variant research.

## Data availability

CobVar database is publicly available at https://datatools.sjri.res.in/VBG/.

## Author contributions

AS and NHS carried out this project and wrote the manuscript. AS was responsible for the conception and design of the project. Data curation were primarily carried out by CAK, AAK, and NHS. NHS took primary responsibility for database and web interface development. AA and TT reviewed the database and implemented the public server setup. CAK, AAK, and NHS reviewed and tested the content and functionality of the online web database. NHS also performed the data analysis for the manuscript. AVK and TT critically reviewed the manuscript, provided feedback and approved the manuscript.

## Acknowledgements

We thank Vaishnavi Chevvu for her assistance in literature review and data curation for the database.

## References

1. Nielsen, M.J., Rasmussen, M.R., Andersen, C.B.F., et al. (2012) Vitamin B12 transport from food to the body’s cells--a sophisticated, multistep pathway. Nat. Rev. Gastroenterol. Hepatol., 9, 345–354.

2. Green, R. (2017) Vitamin B12 deficiency from the perspective of a practicing hematologist. Blood, 129, 2603–2611.

3. Gibson, R. (2005) Principles of nutritional assessment. Principles of nutritional assessment; (2005).

4. Shipton, M.J. and Thachil, J. (2015) Vitamin B12 deficiency - A 21st century perspective. Clin. Med., 15, 145–150.

5. Yetley, E.A., Pfeiffer, C.M., Phinney, K.W., et al. (2011) Biomarkers of vitamin B-12 status in NHANES: a roundtable summary. Am. J. Clin. Nutr., 94.

6. Hunt, A., Harrington, D. and Robinson, S. (2014) Vitamin B12 deficiency. BMJ, 349.

7. Nilsson, S.E., Read, S., Berg, S., et al. (2009) Heritabilities for fifteen routine biochemical values: findings in 215 Swedish twin pairs 82 years of age or older. Scand. J. Clin. Lab. Invest., 69, 562–569.

8. Afman, L.A., Lievers, K.J.A., van der Put, N.M.J., et al. (2002) Single nucleotide polymorphisms in the transcobalamin gene: relationship with transcobalamin concentrations and risk for neural tube defects. Eur. J. Hum. Genet., 10, 433–438.

9. Bailey, L.B. and Gregory, J.F. (1999) Polymorphisms of methylenetetrahydrofolate reductase and other enzymes: metabolic significance, risks and impact on folate requirement. J. Nutr., 129, 919–922.

10. Zinck, J.W.R., De Groh, M. and MacFarlane, A.J. (2015) Genetic modifiers of folate, vitamin B-12, and homocysteine status in a cross-sectional study of the Canadian population. Am. J. Clin. Nutr., 101, 1295–1304.

11. Surendran, S., Adaikalakoteswari, A., Saravanan, P., et al. (2018) An update on vitamin B12-related gene polymorphisms and B12 status. Genes Nutr., 13.

